# Kremen1 dependence receptor induces SEC24C and ATG9A-dependent autophagic cell death

**DOI:** 10.1101/2025.01.15.633131

**Authors:** Sonia Brahim, Thomas Schott, Shiva Ghasemi, Ana Negulescu, Clara Geneste, Elisabeth Errazuriz-Cerda, Gabriel Ichim, Patrick Mehlen, Olivier Meurette

## Abstract

Dependence receptors (DRs) induce cell death by apoptosis when unbound by their cognate ligands. Among them, Kremen1 was first described to induce cancer cell death in the absence of its ligand, DKK1. However, the precise mechanism of Kremen1-induced cell death remains unclear. In this study, we demonstrate that Kremen1 induces cell death with autophagic features, contrasting with the apoptotic process typically associated with dependence receptors. Specifically, the pharmacological inhibition of autophagy, or genetic silencing of key autophagy effectors, efficiently suppresses this cell death process. A biotin proximity labeling for protein-protein interactions identified SEC24C, a component of the COP-II complex, as a critical effector in Kremen1-induced autophagy and cell death. Our findings further reveal that Kremen1 is in proximity with SEC24C and ATG9A after vesicular trafficking and fosters the interaction of SEC24C with ATG8, ERGIC and ATG9A. This potentially underlies the increased number of autophagosomes leading to cell death. The induction of aberrant autophagy by Kremen1 deserves particular attention, especially as the Kremen1/DKK1 pair is frequently altered in cancers. Thus, targeting this pathway may offer a potential strategy for treating cancers resistant to current therapies.

## INTRODUCTION

Dependence receptors (DR) form a family of transmembrane receptor proteins crucial in maintaining cellular homeostasis. Notably, they play a significant role in cancer progression by regulating cell proliferation when bound to their ligand, and by inducing apoptosis in absence of their ligand ^1^. Despite their structural diversity, these receptors constitute a functional family, each shown to induce apoptosis through different mechanisms ^2–4^. Kremen1, a recently identified DR, has been recognized for its ability to induce cell death in absence of its ligand, Dickkopf-Wnt Signaling Pathway Inhibitor (DKK1)^5^. Initially characterized as a Wnt signaling inhibitor, Kremen1/2 serves as a high-affinity receptor for DKK1. Together, they cooperate to recruit the WNT co-receptor LRP5/6, thereby inhibiting the Wnt/β-catenin pathway ^6^. Further studies suggested that Kremen1/2 are not directly involved in this inhibition but they recruit DKK1 and present it to LRP5/6 when the WNT coreceptor is in excess ^7^. Once the ternary complex between DKK1, Kremen1/2, and LRP5/6 is formed, it undergoes rapid clathrin-mediated endocytosis, thereby limiting the number of available Wnt co-receptors on the plasma membrane ^8,9^. Kremen1/2 has been implicated in essential developmental processes, including anteroposterior patterning of the central nervous system (CNS) in *Xenopus* ^10^, as well as DKK1 in mice ^11^. Kremen1 has also more recently been described as a receptor for enteroviruses^12^. Similar to other dependence receptors, the Kremen1/DKK1 receptor-ligand pair may play a significant role in regulating cancer initiation and progression. Kremen1 expression is frequently lost in various cancers, including glioblastoma, neuroblastoma, lung cancer, kidney cancer, stomach cancer, melanoma, and lymphoma ^13^. Conversely, its ligand DKK1 is upregulated in several primary tumor type and is elevated in the serum of cancer patients ^14^. Furthermore, high serum levels of DKK1 are associated with poor prognosis in breast, prostate and bladder cancers ^15–17^, as its overexpression similarly correlated with poor outcomes in hepatocellular carcinoma ^18^. The cell death function of Kremen1 in absence of DKK1 may therefore be implicated in cancer progression. Dependence receptors have been shown to induce apoptotic cell death. However, the mechanism by which Kremen1 induces cell death has not been characterized. We bring here novel insight on this cell death pathway, demonstrating that differently from other dependence receptors Kremen1 is inducing autophagy-dependent cell death in breast and pancreatic cancer cells. This is of particular significance as apoptosis induction may not always be a good strategy in cancer treatment ^19^. We further describe a mechanism involving SEC24C and ATG9A interacting with Kremen1 after its membrane trafficking that fosters their interaction with ATG8, increasing autophagosomes biogenesis.

## RESULTS

### The expression of both Kremen1 and DKK1 pair is frequently altered in cancer

Given the established role of DR and their ligands in tumor suppression, we first examined the expression profiles of the Kremen1/DKK1 pair across various datasets. Consistent with characteristics of tumor suppressors, Kremen1 expression was reduced in multiple cancer types, including breast cancer, colon cancer, and head and neck cancers (Figure 1A). In contrast, DKK1 expression was markedly upregulated across several cancers (Figure 1B). Consistently, analysis of The Cancer Genome Atlas (TCGA) dataset showed that patients with high DKK1 expression had worse overall survival in pancreatic ductal adenocarcinoma and in breast cancers (Figure 1C-D). Interestingly, DKK1 expression is higher in basal-like (TCGA data) and HER2-overexpressing breast cancers (Figure 1E), an observation corroborated in the METABRIC dataset, where DKK1 was elevated in HER2+ and HER2-/ER-subtypes (Figure 1F). In contrast, Kremen1 expression did not vary significantly by subtype in the TCGA dataset, except for a significant difference between luminal B and HER2-enriched (Figure 1G), and a slight but significantly increase in HER2+ cases in the METABRIC dataset (Figure 1H). These findings suggest that alterations in the Kremen1/DKK1 pair may be particularly relevant in basal-like (ER-/HER2-) breast cancers, where *dkk1* gene expression is high while *Kremen1* expression is low. Consequently, we decided to focus on triple-negative breast cancers for further investigation. We also studied pancreatic cell lines given the predictive value of DKK1 in PDAC. Given Kremen1’s role in Wnt inhibition in the presence of DKK1, we examined whether Wnt gene signatures were altered in patients with high Kremen1/DKK1 expression, but observed no significant differences meaning the expression of DKK1 and Kremen1 is not associated to Wnt signature (Figure 1I). Thus, the deregulated Kremen1/DKK1 expression ratio favoring DKK1 is a potential biomarker of poor prognosis.

**Figure 1.**
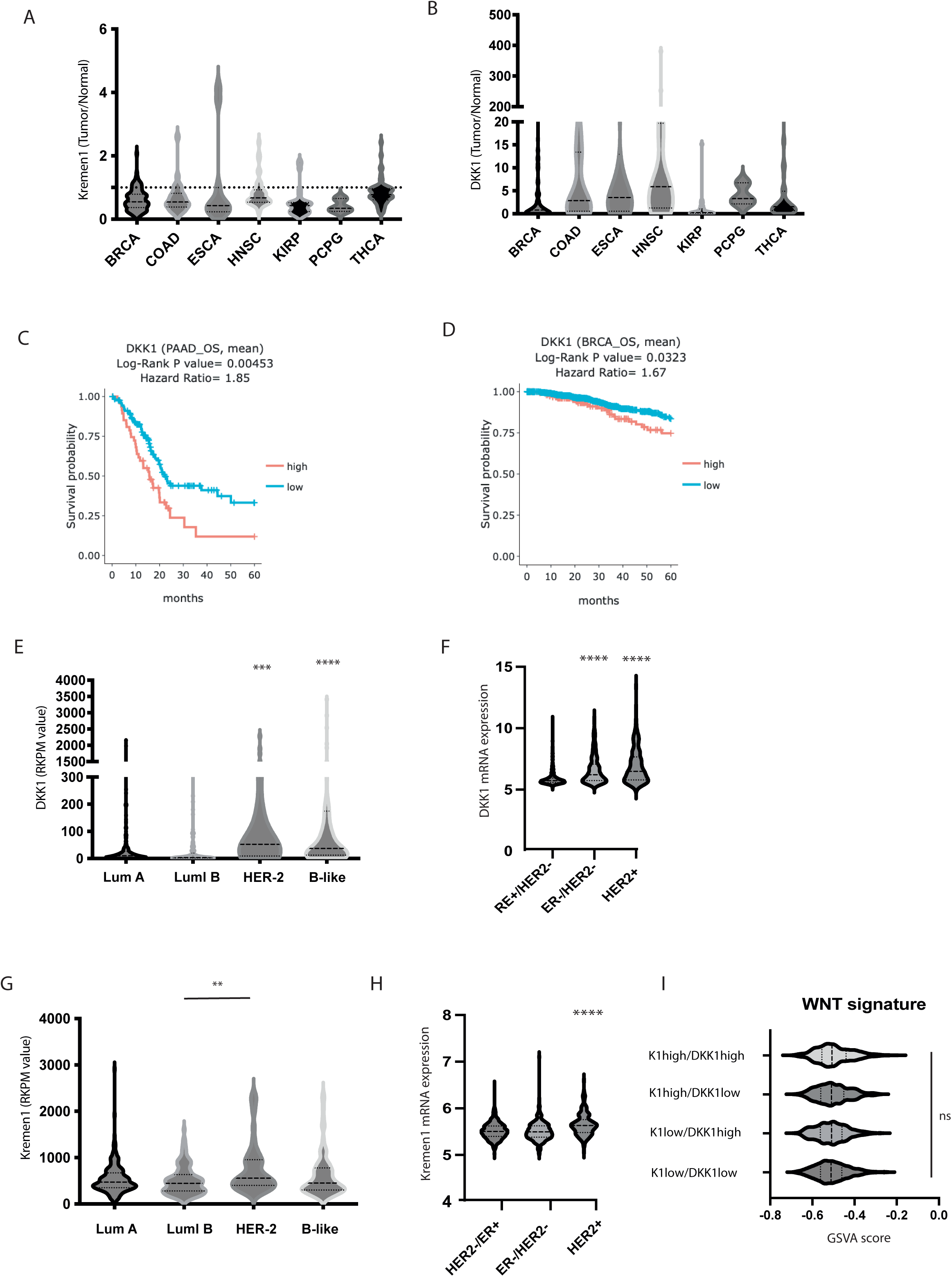
The Kremen-1/DKK1 pair is altered in cancers. (**A,B**) *Kremen1* (A) and *DKK1* (B) expression was assessed in paired Normal /Tumoral sample from different cancers from TCGA dataset BRCA: Breast invasive carcinoma; COAD: Colon adenocarcinoma; ESCA: Esophageal carcinoma; HNSC: Head and Neck squamous cell carcinoma; KIRP: Kidney renal clear cell carcinoma; PCPG: Pheochromocytoma and Paraganglioma; THCA: Thyroid carcinoma. (**C,D**) Kaplan-Meier survival analysis of *DKK1* expression (low or high) in pancreatic adenocarcinoma (**C**) and breast cancer (**D**) patients from the TCGA dataset. (**E-H**) Violin plot of *DKK1* (**E,F**) and *Kremen1* (**G,H**) expression across different subtypes of breast cancer patients from the TCGA (**E,G**) and metabric datasets (**F,H**). One-way ANOVA was performed with a Tukey’s multiple comparison test (*** p-value < 0.001; **** p-value < 0.0001).

### Kremen1-induced cell death lacks classical apoptotic features

Kremen1 has previously been described as a DR inducing features of apoptotic cell death^5^. In order to investigate the mechanisms underlying Kremen1-induced cell death, we developed several stable cell lines expressing Kremen1 upon doxycycline treatment. Kremen1 was induced in a time and dose dependent manner (supplementary figure 1A). Furthermore, whereas doxycycline treatment had no impact on cell death (supplementary Figure 1B), Kremen1 induction was followed by a significant increase of cell death: up to 50% in MDAMB231 cells, 30% in HEK293 and 20-30% in B<u>x</u>PC3 (supplementary figure 1C-E), showing that Kremen1-induced pathway is conserved across different cell types. We further focused on MDAMB231 and BXPC3, as model of <u>b</u>reast cancer and pancreatic cancer and used IncuCyte Imager-based live cell imaging to monitor cell death every 2 hours upon a long period of time. We observed both in MDAMB231 and BxPC3 accumulation of cell death upon Kremen1 induction from 24 up to 60 hours following Kremen1 induction (Figure 2A, supplementary Figure 1F). Additionally, long-term clonogenic survival revealed a reduction in colony formation upon Kremen1 induction (Figure 2B). Co-expression of DKK1 reduced Kremen1-induced cell death, supporting the previously documented DR activity of Kremen1 (Figure 2C). Furthermore, siRNA targeting DKK1 also induced cell death although to a lower extend than Kremen1 induction (Supplementary Figure 1G). Kremen1 expression led to activation of caspase-3, which was efficiently inhibited by treatment with Q-VD-OPh pan-caspase inhibitor (supplementary Figure 2A). However, treatment with Q-VD-OPh (QVD) inhibited only very partially cell death in MDAMB231 (Figure 2D) and had no effect in HEK293 cells either (Supplementary figure 1D). We also treated cells with additional caspase inhibitors, including IETD and DEVD, and observed no impact on cell death, indicating that these caspases are not involved in our cell death model (data not shown). Interestingly, treatment of Kremen1-expressing cells with ABT263 triggered a rapid shift towards apoptotic features, indicating that these cells remain responsive to apoptotic stimuli and that Kremen1-induced caspase activity may prime cells to a stronger apoptotic stimulus (Supplementary Figure 2B). We further characterized cell morphology by Nanolive microscope-based holotomography imaging of cells expressing Kremen1 and of cells treated with an apoptosis inducer as a control (BV6). Cells treated with BV6 exhibited membrane blebbing and cell contraction, while Kremen1-overexpressing cells displayed a phenotype characterized by cellular swelling (Figure 2E). Treating cells with necrostatin-1 or ferrostatin did not rescue cell death induction showing that Kremen1 is not inducing necroptosis or ferroptosis (Figure 2F and supplementary figure 2C). Thus, our results indicate that kremen1 induces a form of cell death distinct from classical forms of programmed cell death.

**Figure 2:**
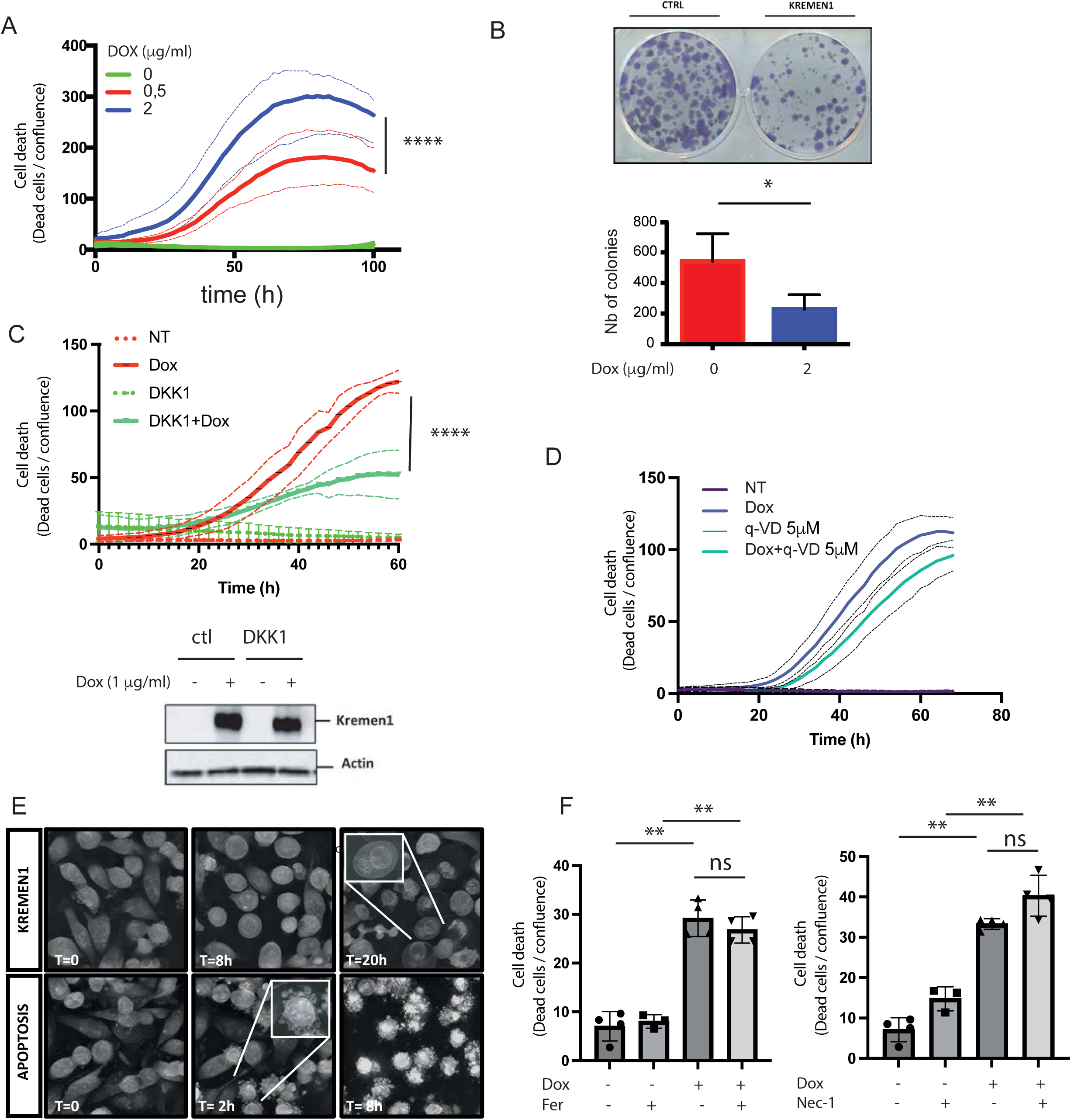
Kremen1 does not induce apoptotic cell death (**A**) Cell death was measured by propidium iodide incorporation in MDA-MB-231 with or without Kremen1 expression upon doxycycline treatment (0,5 and 2 mg/mL). Two-way ANOVA was used to assess effect of concentration on cell death induction (****: p-value <0.0001). (**B**) Colony formation assay using crystal violet with or without Kremen1 expression following dox treatment (dox 2mg/mL) and quantification of the colonies. Student’s t-test was performed to compare mean of two groups. (*: p-value<0.05). (**C**) Cell death measured by propidium iodide incorporation upon Kremen1 expression in presence or not of DKK1 expression and western blot analysis showing the expression of Kremen1 is similar in the two cell lines. Two-way ANOVA test was used to compare control and DKK1 expressing groups. (****: p-value<0.0001). (**D**) Cell death was measured by propidium iodide incorporation every two hours with the incucyte after treatment with Doxycycline (1mg/ml) to induce Kremen1 expression in presence or not of qVD-oph treatment (5mM). (**E**) Nanolive imaging analysis comparing Kremen1 expressing cells and apoptotic cells treated with BV6 (1mM). (**F**). Cell death of MDAMB231 cells 30 hours after treatment with doxycycline (1mg/ml) to induce Kremen1 expression in presence or not of Ferrostatin (Fer, 10mM) or necrostatin-1 (Nec, 10mM). One-way ANOVA was performed with Tukey’s multiple comparison test. (**: p-value<0.01).

### Kremen1 induced-cell death displays autophagic features

Prior to examining the specific mechanisms of Kremen1-induced cell death, we analyzed the morphology of dying cells using transmission electron microscopy (TEM). We did not observe apoptotic features like membrane blebbing or nuclear condensation and cells appeared to increase in volume rather than exhibit the typical apoptotic shrinkage (Figure 3A). Across multiple time points, in both adherent and non-adherent cells, the dying cells consistently displayed autophagic features, including an accumulation of vacuoles, phagophores, autophagosomes, and multilamellar bodies (supplementary figure 3A). In the supernatant, rare apoptotic cells were observed although most of dead cells appeared necrotic (Supplementary figure 3B). We confirmed autophagy induction through molecular analyses, evidenced by the accumulation of the lipidated form of LC3 and a decrease in p62 expression upon Kremen1 induction, as shown by western blotting (Figure 3B), and the formation of dense LC3 puncta in immunofluorescence assays (Supplementary Figure 4A,B) indicating that autophagosomes are increased in dying cells both in MDA-MB231 and BxPC3 cells. Moreover, no LC3 lipidation was observed in control cells treated with doxycycline (Supplementary figure 4C). In agreement with autophagy being regulated by the DR function of Kremen1, siRNA targeting DKK1 induced autophagic features as does Kremen1 expression (supplementary figure 4D). We next characterized autophagic flux by expressing Kremen1 in cells expressing the RFP-GFP-LC3 reporter construct. We observed accumulation of RFP-only LC3 dots upon Kremen1 induction demonstrated an induction of the autophagic flux and not a mere inhibition of late autophagic phase leading to accumulation of autophagosomes (Figure 3C).

**Figure 3:**
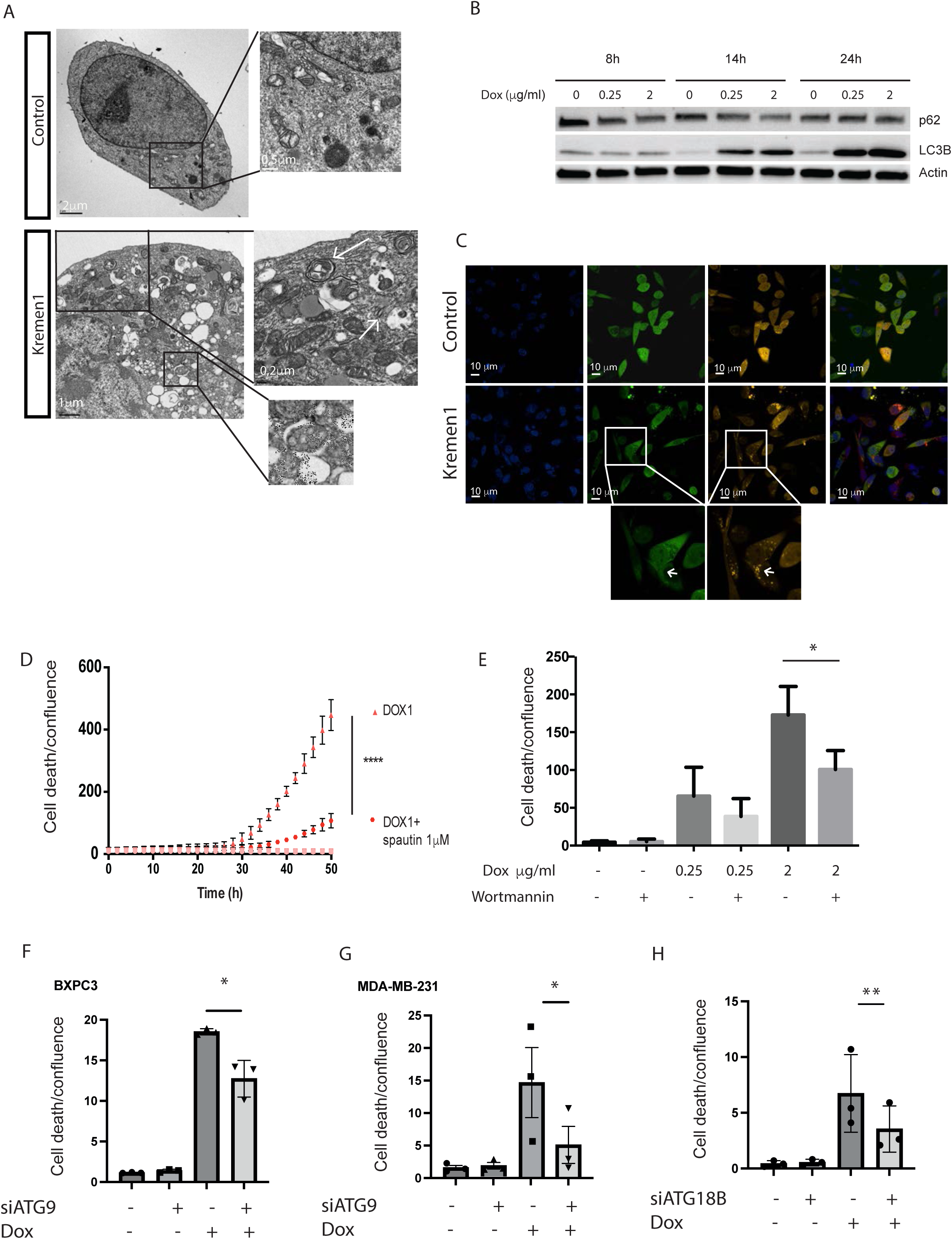
Kremen1-induced cell death displays autophagic features (**A**) Transmission electronic microscopy of MD-AMB-231 cells expressing Kremen1 showing numerous vacuoles and autophagic features (white arrow). (**B**) Western blot showing LC3 and p62 expression upon doxycycline treatment with the indicated concentrations at 8h, 14h and 24 hours. (**C**) Confocal images of MDAMB231 cells expressing the RFP-GFP-LC3 reporter treated with doxycycline (Kremen1) or not (Control) for 12 hours. (**D**) Measure of cell death by propidium iodide incorporation following Kremen1 expression (DOX) in presence or not of Spautin-1 (1mM) (**E**) Measure of cell death by propidium iodide incorporation following Kremen1 expression (DOX) in presence or not of Wortmanin (5mM). (**F-G**) Monitoring of cell death following transfection with siRNA targeting ATG9A in BxPC3 cells (**F**) or MDAMB231 (**G**) followed by Kremen1 induction (Dox; 1 mg/ml). (**H**) Monitoring of cell death of MDA-MB-231 cells treated (Dox; 1 mg/ml) or not to induce Kremen1 following transfection with siRNA targeting ATG18B . Ratio paired Student’s t-test was used to test the effect of knock-down on cell death on three independent experiment (*: p-value<0.05 ; **: p-value<0.01).

### The autophagic machinery is indispensable for Kremen1-induced cell death

We next sought to assess the requirement of the autophagic machinery in the observed cell death. Spautin as well as wortmanin treatment which inhibit the early stage of autophagic flux by preventing autophagosome formation, resulted in a significant decrease in cell death and LC3 lipidation (Figure 3 D,E, supplementary figure 5A,B). In contrast, treatment with Bafilomycin-A and Chloroquine, which both block the flux at late stages inducing autophagosome accumulation led to an increase of cell death following kremen1 expression (Supplementary Figure 5C). This suggests that the presence of autophagosomes or certain autophagic components may be essential for cell death induction. While autophagy is often induced following apoptosis as a mechanism thought to inhibit cell death, in our model, treatment with Spautin1 completely blocked cell death by inhibiting LC3 lipidation (Figure 3D, supplementary Figure 5B) without affecting caspase activation (Supplementary figure 5D). Furthermore, knock-out of APAF1 which inhibits caspase activation had no effect on cell death nor on LC3 lipidation (supplementary Figure 5E-G), suggesting that caspase activation in not involved in autophagy induction in this context. Surprisingly, knock-down of ATG5 by siRNA or by CRISPR knock-down had no impact on cell death induction nor on caspases activation (data not shown) demonstrating the involvement of atypical autophagy. As ATG5 was not involved in Kremen1-induced cell death, we examined the role of ATG9A, since it is a protein which facilitates membrane recruitment and autophagosome membrane expansion ^20^. We further showed that inhibition of ATG9 via siRNA (supplementary Figure 6A) significantly reduced cell death in both MDA-MB-231 and BxPC3 cell lines, highlighting its essential role in this process (Figure 3F-G). We also investigated the involvement of ATG18 since they have shown to participate with ATG9A to autophagosome membrane expansion ^21^. siRNA mediated knock-down of ATG18B also partially rescued Kremen1-induced cell death (Figure 3H). Altogether, these findings support the involvement of an atypical form of autophagy relying on ATG9A and ATG18B to induce aberrant accumulation of autophagosomes leading to autophagic cell death.

### SEC-24C is involved in Kremen1-induced cell death

To elucidate the mechanisms underlying this autophagy-mediated cell death, we analyzed the Kremen1 interactome via mass spectrometry. For this purpose, we employed a biotin proximity labeling for protein-protein interactions proximity labeling approach by fusing Kremen1 with an ascorbate peroxidase (APEX2) enzyme to perform enzyme-mediated biotin identification. Using the same doxycycline-inducible system, we expressed a Kremen1-APEX fusion protein to identify proteins in close proximity. We first confirmed that the Kremen1-APEX fusion did not alter Kremen1-mediated signaling, and induced autophagy and cell death with comparable efficiency to Kremen1 alone (data not shown). We then used streptavidin beads to isolate proteins proximal to Kremen1-APEX, followed by mass spectrometry analysis, which yielded a list of proteins most enriched upon doxycycline addition and subsequent Kremen1 induction (Figure 4A,B). Among the most enriched proteins were SEC24C and other proteins of the COP-II complex. Two major clusters appeared using String-database analysis (Figure 4B). The first one was enriched in membrane-associated, adhesion and focal adhesion proteins such as integrins (ITGB1, ITGA6, ITGA2), taline (TLN1), vinculin (VCL), paxillin (PXN), showing the membrane localization of Kremen1 in the experiments. The second cluster was a cluster of COP-II complex proteins, including SEC23A and B, SEC24 B and C as well as SEC31A. This complex particularly caught our attention since the COP-II complex has been implicated in atypical autophagy regulation ^22,23^. Specifically, SEC24C stands out due to its role in autophagy regulation by its interaction with ATG9 to increase autophagosomes abundance in yeast and is implicated in STING-induced autophagy and cell death ^24,25^. Immunofluorescence analysis confirmed partial interaction between Kremen1 and SEC24C (Figure 4C). To investigate the potential role of SEC24C during Kremen1 cell death induction, we inhibited its expression via siRNA and found a significant reduction of Kremen1-induced cell death in both MDA-MB-231 and BxPC3 cell lines (Figure 4D). Furthermore, knocking-down of SEC24C induced a significant reduction of LC3 lipidation upon Kremen1 induction (Figure 4E). Altogether these data demonstrate that SEC24C is involved in Kremen1-induced autophagy and cell death.

**Figure 4:**
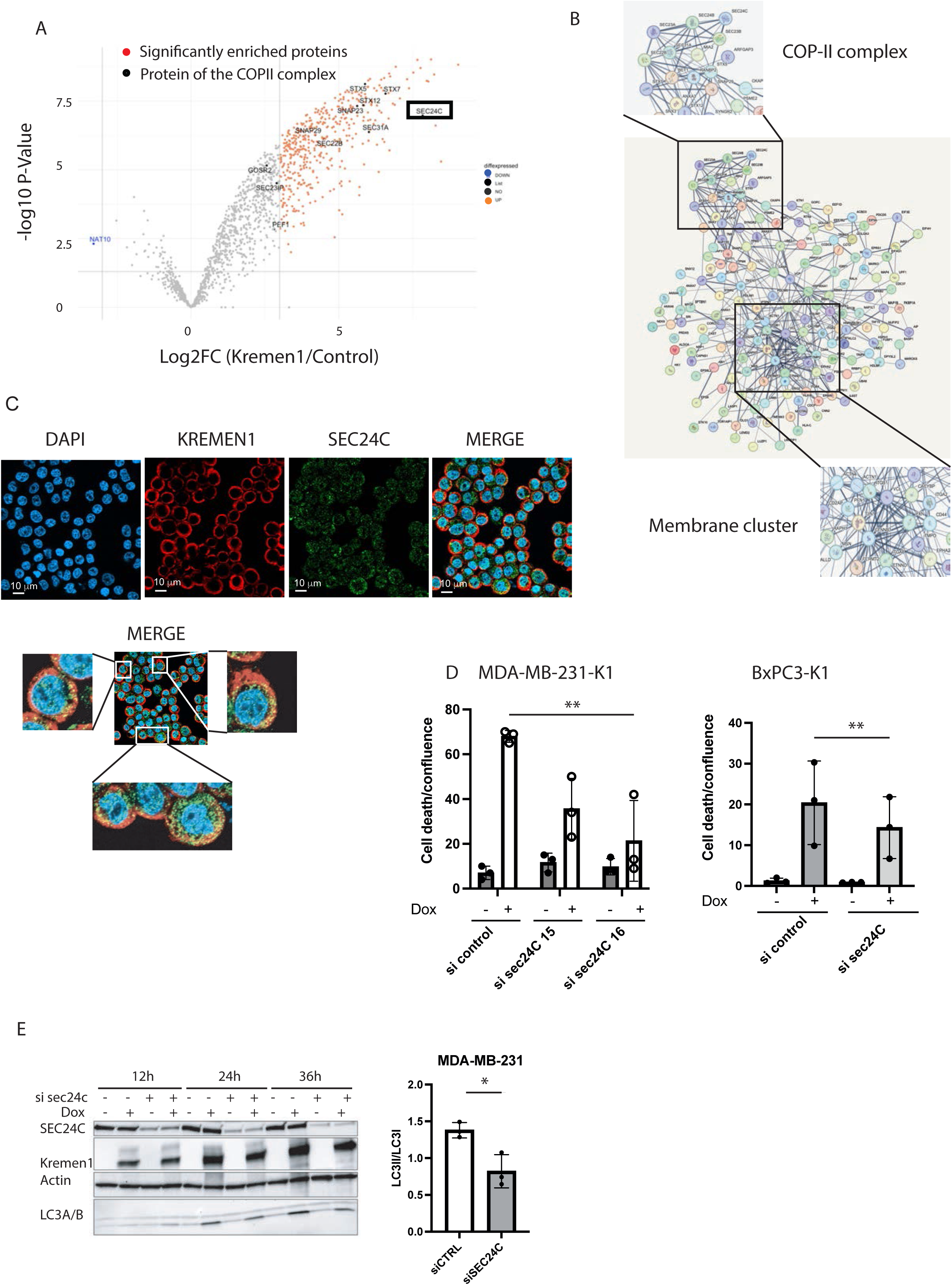
SEC24C is implicated in Kremen1-induced cell death (**A**) Volcano plot showing protein identified by mass spectrometry in Kremen1 interactome. (**B**) STRING database protein interaction network map highlighting members of the COP-II complex revealed by mass spectrometry. (**C**) Confocal imaging showing cells stained with DAPI, Kremen1 antibody (red) or Sec24C antibody (Green) and merged on BxPC3 cell line treated with doxycycline (1 mg/ml) to induce Kremen1. (**D**) Cell death induction following Kremen1 induction (dox, 2mg/ml) and knock-down of SEC24C by siRNA in MDA-MB -and BxPC3 cells. Ratio paired Student’s t-test was used to test the effect of knock-down on cell death on three independent experiment (**: p-value<0.01). (**E**) Western blot analysis of SEC24C, Kremen1, Actin and LC3 expression following inhibition of Sec24C by siRNA at 12h, 14h and 36h and associated quantification at 24h.

### Kremen1 is at proximity of SEC24C and ATG9 after being internalization

As SEC24C and the COPII complex are involved in protein trafficking, we further wanted to characterize the localization of SEC24C and ATG9 at proximity of Kremen1 after membrane trafficking. We set up Proximity labelling assay (PLA) by using Kremen1/SEC24C and Kremen1/ATG9A antibodies and could observe significant co-localization of Kremen1 with ATG9A and SEC24C spread in the cytoplasm (Figure 5A). Furthermore, DKK1 expression together with Kremen1 induces a reduction of Kremen1 interaction with ATG9A (Supplementary figure 6B), suggesting that Kremen1 has to be addressed to the membrane to further interact with ATG9A. To test this hypothesis, we performed pulse chase experiment by adding Kremen1 antibody on live, non-permeabilized cells and performed PLA with adding ATG9A or SEC24C antibodies after different time points. By this technique we visualize the proximity of internalized Kremen1 (iK1) with intracellular SEC24C or ATG9A. We could demonstrate a time dependent increase in PLA signals for internalized Kremen1/SEC24C and internalized Kremen1/ATG9A. Altogether, these experiment<u>s</u> show that localization of Kremen1 with SEC24C and ATG9A is at least in part dependent on Kremen1 addressing to the membrane and is dependent on DKK1.

**Figure 5:**
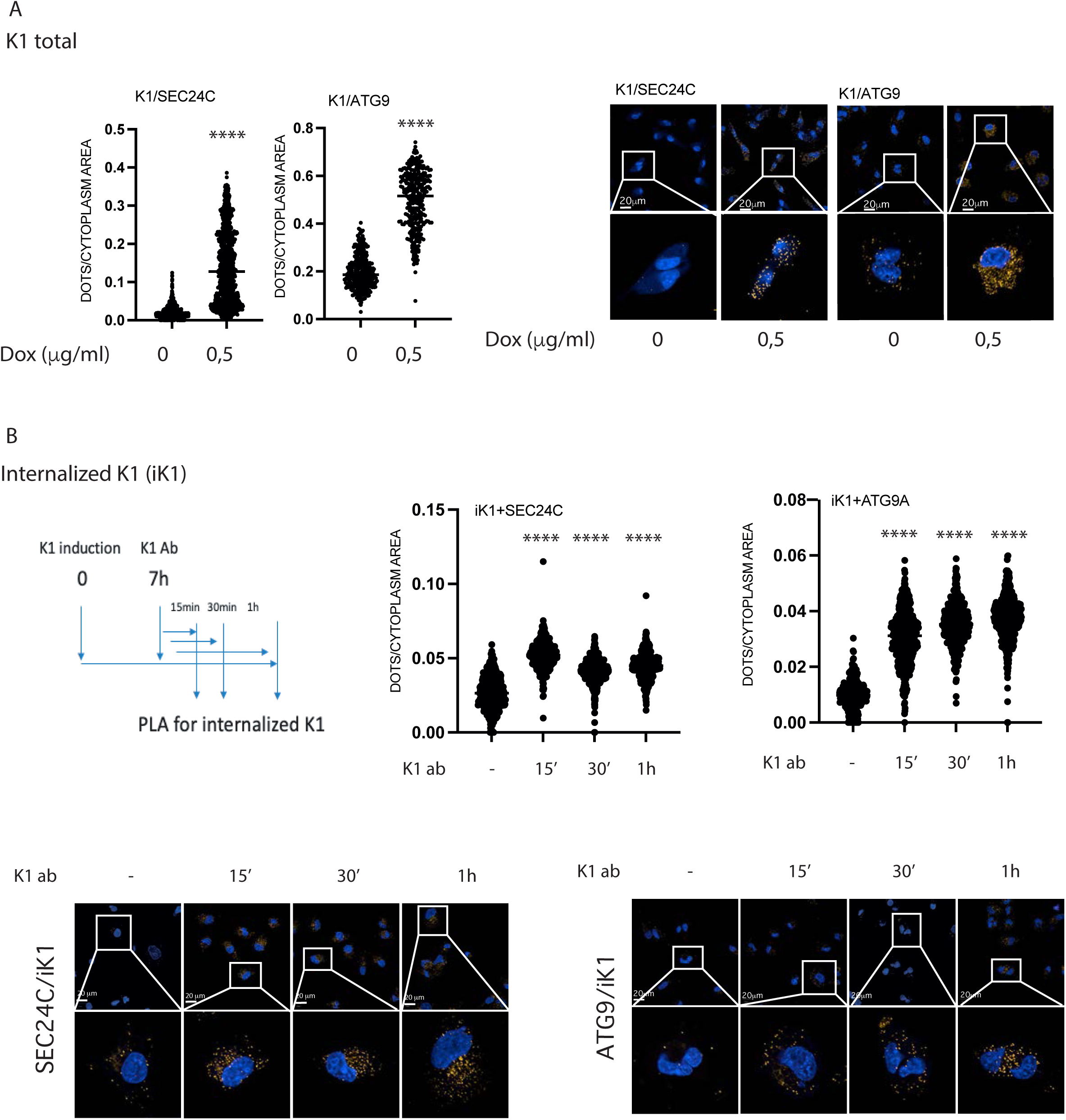
Kremen1 is interacting with ATG9A and SEC24C after membrane trafficking (**A**) PLA images and quantification of Kremen1/SEC24C and Kremen1/ATG9A interaction following Doxycycline treatment (0,5 mg/ml) to induce Kremen1. Student’s t-test was performed on automated counting using Signal Image Artist software (****: p-value< 0,0001). (**B**) PLA was performed after incubation of live cells with Kremen1 antibody as indicated in the scheme. PLA was then performed with only SEC24C or ATG9 antibody to test proximity with internalized Kremen1 (iK1). Student’s t-test was performed on automated counting using Signal Image Artist software (****: p-value< 0,0001).

### Kremen1 induces recruitment of ATG8 at ERGIC (LMAN1) together with SEC24C and ATG9A

As we observed that SEC24C was involved in Kremen1 induced autophagy, we then sought to determine whether SEC24C was associated with autophagosomes upon Kremen1-dependent cell death induction. To this end, we performed immunoprecipitation of SEC24C in the presence and absence of Kremen1 and observed that, upon Kremen1 induction, SEC24C interacted with ATG8, indicating that SEC24C plays a specific role in autophagy regulation within this autophagic cell death pathway (Figure 6A). This observation was further validated through confocal microscopy, which revealed a partial colocalization of ATG8 and SEC24C in BxPC3 cells in condition of Kremen1 expression, as indicated by yellow puncta (Figure 6B). We next demonstrated that upon Kremen1 induction, ATG8 localization at ERGIC (LMAN1) together with SEC24C and ATG9 was increased (Figure 6C). Altogether, these findings demonstrate that SEC24C and ATG9 are essential components in mediating Kremen1-induced cell death by increasing autophagosome biogenesis.

**Figure 6:**
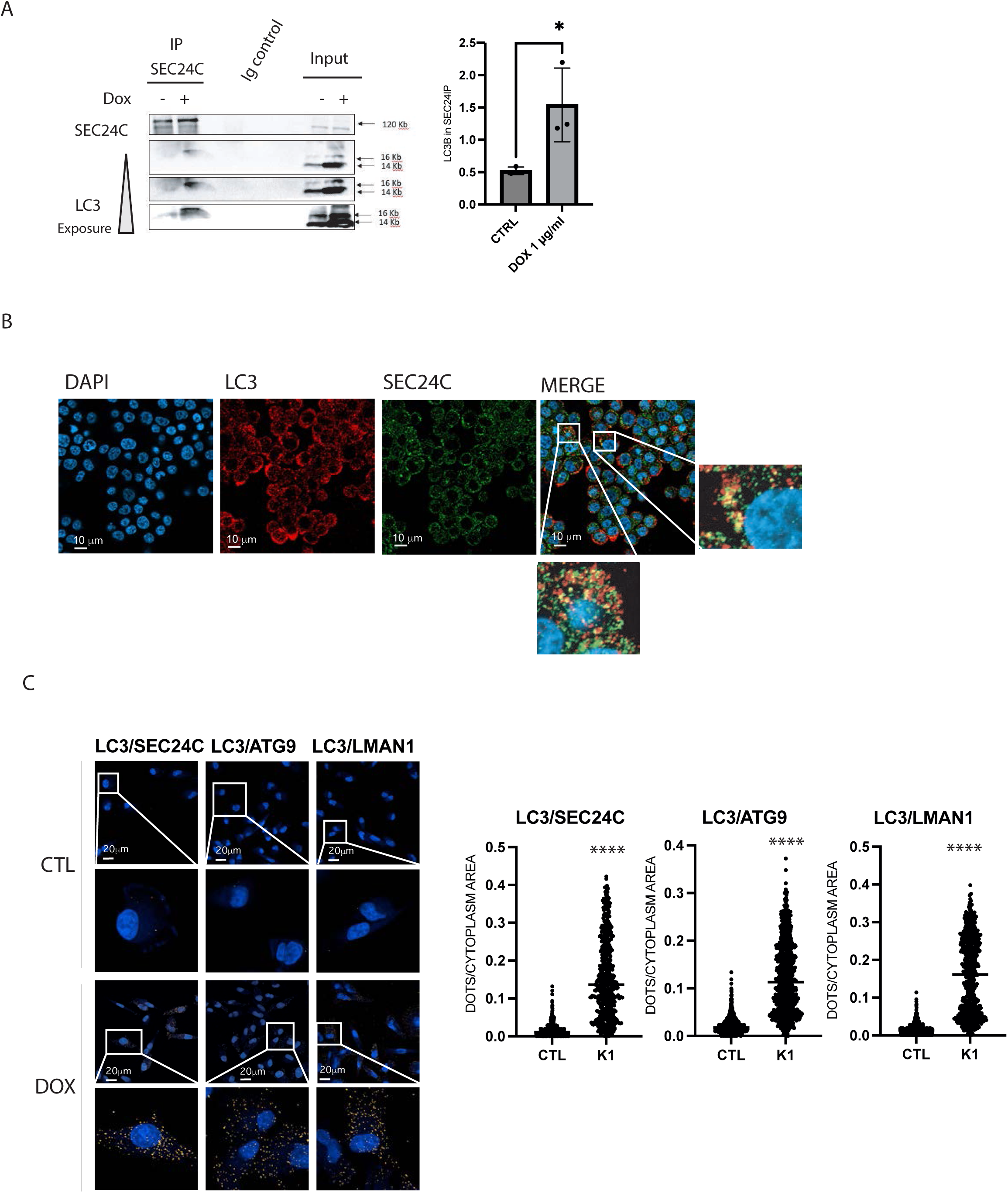
Kremen1 induces SEC24C/ATG9A/ATG8 interaction at ERGIC (**A**) Western blot analysis following immunoprecipitation of SEC24C upon Kremen1 expression and associated quantification. (**B**) Confocal imaging showing cells stained with LC3 (red) SEC24C (Green) or DAPI following Kremen1 induction. (**C**) PLA images and quantification of LC3/SEC24C, LC3/ATG9A and LC3/ERGIC (LMAN1) interaction in presence of Kremen1 induction Student’s t-test was performed on automated counting using Signal Image Artist software (****: p-value< 0,0001).

## DISCUSSION

Kremen1, initially identified as a receptor involved in DKK1-mediated Wnt signaling inhibition ^6^, was later characterized as a DR ^5^. DRs constitute a diverse family of structurally unrelated receptors that share the unique capacity to actively induce cell death in the absence of their respective ligands. Until now, all known DRs have been shown to induce apoptosis through distinct pathways that ultimately converge to caspase-3 activation ^26^. In the case of Kremen1, Causeret et al. reported that its overexpression led to caspase-3 cleavage, consistent with an apoptotic pathway similar to that of other DRs, but without any characterization of the cell death signaling pathway. In this study, we confirmed that Kremen1 expression induces caspase-3 activation despite the absence of classical apoptotic features, as evidenced by transmission electron microscopy and time-lapse tomographic microscopy. Moreover, inhibition of caspase activity <u>with</u> qVD-Oph had very little effect on cell death indicating that effector caspases are dispensable for Kremen-1 induced cell death. It is plausible to consider that caspase activation in this context occurs as a bystander event, insufficient to directly induce cell death but potentially involved in non-apoptotic roles as it already has been shown before ^27^. Another explanation would be that the aberrant increase of autophagosomes may prevent the cleavage of canonical substrates necessary for apoptotic progression. However, cells expressing Kremen1 remain responsive to apoptotic stimuli, particularly in the early stages following Kremen1 induction, as treatment with ABT263, a Bcl-xL and Bcl-2 antagonist, rapidly induces canonical apoptosis indicating that cells remain sensitive to apoptotic stimuli.

Instead of apoptotic features, we consistently observed autophagic features in dying cells. Cells exhibiting autophagic features have long been reported in physiological cell death ^28^. However, the presence of abundant autophagosomes in dying cells does not necessarily imply that autophagy is the direct trigger of cell death. Autophagy is well-established as a survival pathway under nutrient-limiting conditions ^29^ and it is generally considered to limit apoptosis and confer resistance to cell death ^30^. Evidence from physiological development supports the role of autophagy as a *bona fide* cell death mechanism. In Drosophila, for example, autophagy, not apoptosis, is essential for programmed cell death in the midgut ^31^ and is also required for the degradation of salivary gland cells ^32^. Additionally, autophagy-dependent cell death, termed autosis, can be induced by autophagy-stimulating peptides, such as Tat-Beclin1, which holds potential for therapeutic applications ^33^. The cell death observed following Kremen1 induction displays phenotypic similarities with Tat-Beclin1-induced autosis. However, treatment with Na^+^/K^+^ inhibitors did not rescue this cell death (data not shown), suggesting mechanistic differences from canonical autosis. Kremen1-induced cell death results in a substantial accumulation of autophagosomes, which may contribute to the observed cytotoxicity. Autophagosome accumulation alone has been reported as sufficient to induce cytotoxic effects ^34^ and the timing of autophagosome accumulation relative to ATP availability and lysosomal homeostasis may be <u>a</u> critical factors ^35^. Although protein overexpression may induce cellular stress triggering autophagy, the phenotype observed with Kremen1 has never been observed with other DRs using the same induction strategy ^3^. Furthermore, DKK1 addition is reducing cell death in conditions in which Kremen1 expression is induced at the same levels (Figure 1C). By a proximity biotin ligation assay, we identified enrichment of members of the COP-II complex associated with Kremen1 (Figure 4). The COP-II complex primarily serves two functions: it facilitates the physical deformation of the endoplasmic reticulum membrane into vesicles and selects cargo molecules for transport to the Golgi apparatus. Interestingly, COP II vesicle are diverted to autophagosome formation in response to stresses ^36,37^. Furthermore, different members of this complex have been shown to be involved in autophagosomes formation ^38–40^. More specifically, SEC24C, identified as the most enriched component of the COPII complex within the Kremen1 interactome, functions as a pivotal effector in autophagic process, including ER-Phagy ^22^ as well as STING-associated autophagy and cell death ^24,41^. Furthermore, under cellular stress, such as nutrient deprivation, SEC24C undergoes phosphorylation by the kinase Hrr25, enabling it to interact with ATG9 ^38^. Experimental evidences demonstrate that SEC24C and ATG9A are implicated in Kremen1-mediated induction of autophagic cell death. Pulse-chase labeling analyses have further revealed colocalization of Kremen1 with both ATG9A and SEC24C following membrane trafficking. This interaction facilitates LC3 redistribution to ATG9A, the ER-Golgi intermediate compartment (ERGIC) and SEC24C illuminating previously uncharacterized aspects of the Kremen1-initiated signaling cascade. Unlike other DRs Kremen1 orchestrates a unique cell death modality centered on the autophagic machinery. We also showed that in presence of its ligands DKK1, Kremen1 is not or much less interacting with ATG9A. We therefore hypothesize that upon Kremen1 induction or limitation for DKK1 ligand, Kremen1 is recruiting ATG9A vesicle to SEC24C/ATG8 site of autophagosome formation creating an increased demand for membrane resources to support autophagosome formation.

Given the relevance of DRs as therapeutic targets in cancer ^26^, we examined the status of the Kremen1/DKK1 pair across various cancer types. Our observations indicate that Kremen1 expression is frequently downregulated in multiple cancers, suggesting a potential role for Kremen1 as a tumor suppressor through its autophagy-dependent cell death pathway. Notably Sato et al ^14^ reported elevated DKK1 serum levels in patients, stronger than those observed here in RNA sequencing data. In breast cancer specifically, while some patients exhibited high DKK1 levels, the median expression of DKK1 in matched tumor/normal samples remained below 1 in breast tumors (Figure 1B), contrasting with the significant serum increases observed by Sato et al,. Additionally, DKK1 expression has been linked to metastasis and poor survival outcomes in lymph node-positive breast cancer patients ^42^. Notably, recent studies have demonstrated that autophagic cell death may contribute to tumor suppression by limiting chromosomal instability ^43,44^. We are currently exploring the therapeutic potential of activating this novel autophagic cell death pathway as a targeted approach in cancer treatment.

## EXPERIMENTAL PROCEDURES

### Plasmids construction

The inducible Sleeping Beauty transposon system was used as backbone for cloning the murin Kremen1 gene under doxycycline inducible control. The Kremen1 gene was fused in N-terminal with an HA tag. Two constructs were realized: pitr1-Kremen1, containing a GFP insertion constitutively expressed and a puromycin resistant gene constitutively expressed and pitr2-Kremen1 containing a RFP insertion and a zeocin resistant gene both constitutively expressed.

### Cell culture

Human breast cancer cell line MDA-MB231 and human embryonic kidney HEK293T cells were obtained from ATCC (UK) and maintained in Dulbecco’s Modified Eagle’s Medium (DMEM) supplemented with 10% fetal bovine serum (FBS) and 1% penicillin/streptomycin.

For the establishment of stable cell lines MDA-MB231 and HEK293T with pitr1-Kremen1 or pitr2-Kremen1, the cells were transfected with 1,6 μg plasmid, 0,4 μg SB100x transposase and Fugen (Promega) in a ratio total DNA : transfectant of 1:3 and then selected with puromycin (Gibco Life Technologies) or zeocin (Invitrogen) for 2 to 4 weeks. MDA-MB231 clones were selected by flow cytometry in accordance with high or low expression of GFP.

For inhibitor treatments cells were incubated with 20 μM q-VD (Sigma-Aldrich SML0063), 40 μM Necrostatin-1 (Sigma-Aldrich N9037), 5 mM 3-MethylAdenin (Sigma-Aldrich M9281) in different combinations. For induction of Kremen1, Doxycycline (Sigma-Alderich) was added in various concentrations in cell media.

siRNA transfections were realized in 6-well plates with 10nM siRNA and 3μl Lipofectamine RNAiMAX (Life) according to the manufacturer instructions. The siRNA used were Life Silencer Select Negative control #1 Cat#4390843, lot AS021N56;

### Transmission electron microscopy (TEM)

Cells were prepared for fixation in 6-well plates or in supernatants with 1 volume complete DMEM medium/1 volume glutaraldehyde 4% for 15 minutes at 4°C. Cells were fixed in 1 volume cacodylate pH 7,4/1volume g glutaraldehyde 4%. Attached cells were harvested by scratching and centrifugated. The sample were washed three times in saccharose 0.4 M/0.2 M Na C-HCl-Cacodylate-HCl pH 7.4 à 0,2 M for 1 hour at 4°C, and postfixed with 2% OsO4/0.3 M Na C-HCl Cacodylate-HCl pH 7.4 1hour at 4°C. Then cells were dehydrated with an increasing ethanol gradient (5minutes in 30%, 50%, 70%, 95% and 3 times for 10 minutes in absolute ethanol. Impregnation was performed with Epon A (50%) plus Epon B (50%) plus DMP30 (1,7%). Inclusion was obtained by polymerisation at 60°C for 72hrs. Ultrathin sections (approximately 70 nm thick) were cut on a ultracut UC7 (Leica) ultramicrotome, mounted on 200 mesh copper grids coated with 1:1000 polylisine, and stabilized for 1day at room temperature (RT) and, contrasted with uranyl acetate and lead citrate. Sections were examined with a Jeol 1400 JEM (Tokyo,Japan) transmission electron microscope equipped with a Orius 600 camera and Digital Micrograph.

### Caspase-3 activity assay

Cell<u>s</u> were harvested by trypsinization and cell pellets were obtained by centrifugation at 4°C and lysed. The caspase-3 activity assay was performed according to the manufacturer’s instructions (Biovision caspase-3 colorimetric assay kit). Total protein concentrations were measured with the BCA assay kit using BSA as a standard (Pierce Biotechnology, Rockford, IL, USA). Absorbance readings were done on a TECAN infinite F500.

### PI/Sytoxgreen stained cells counting

Cells were plated in 96-well plates, incubated with 0,33 ng/ml PI or Sytoxgreen and different treatments. The plates were further monitored by Incucyte Zoom (Essenbio) for several days. The PI positive cells were normalized by the confluence percentage for each well, at several time points.

### Immunofluorescence staining

Fixation and staining were performed simultaneously. Attached cells samples were fixed with freshly prepared 4% PFA, washed 3 times with PBS for 5 minutes and permeabilized with PBS-0.2% Triton X-100 (TX-100) for 10 minutes at room temperature. Samples were then washed 3 times with PBS for 5 minutes and blocked in PBS with 4% bovine serum albumin (BSA), 2% normal donkey serum and 0.2% TX-100 for one hour. Primary antibodies were diluted in the blocking solution at a dilution of 1:200 for the anti-Kremen1 produced antibodies and 1:50 for anti-HA (Sigma-Aldrich H4908). Alexa-conjugated secondary antibodies (Alexa555-donkey anti rabbit, Alexa488-donkey anti rabbit, Alexa488-donkey anti mouse) were used at 1:1000 dilution. 1 µg/ml DAPI was added at the end to stain nuclei. Images were acquired with Zeiss Axio fluorescence microscopy.

### Quantitative RT PCR

mRNAs were extracted with the NucleoSpin RNA kit (Machery-Nagel) according to manufacturer’s instructions. cDNAs were generated with the iScript cDNA Synthesis kit (BIO-RAD) according to the manufacturer’s instructions. Real-time quantitative RT-PCR was performed using a LightCycler 480 (Roche Applied Science).

### Western Blot

Cells were lysed in SDS buffer (2% SDS, 150mM NaCl, 50mM Tris-HCl, pH 7.4), denaturated at 94 C° for 5 minutes. Protein concentration was measured with the BCA assay kit using BSA as a standard (Pierce Biotechnology, Rockford, IL, USA) according to manufacturer’s instructions. The primary antibodies used were : anti-HA (1:1000 dilution, Sigma-Aldrich H4908), anti-LC3 (1:1000 dilution, Cell signaling ) and an anti-actin conjugated with HRP ().

### Proximity-Dependent Biotinylation assay

In order to identify new partners of the Kremen1 induced signalization, a proximity biotinylation screening of the receptor Kremen1 was performed. The receptor Kremen1 was fused to the soybean ascorbate peroxydase (APEX2) in a Tet–ON/OFF plasmid (PITR). MDA-MB 231 cell line was stably transfected with the PITR Kremen1-APEX plasmid expressing the receptor fused to APEX under DOX treatment. For each biotinylation experiment 30 million cells were seeded. 2 conditions were chosen, a control condition without induction of Kremen1 (DOX 0) and the dox treated condition (DOX 0.25). DOX (0.25 µg/mL) was added 24 hours after seeding and biotinylation was performed after 24 hours of treatment. 500 µm Biotin Phenol (BP) in DMEM and incubated for 30 minutes at 37°C. Fresh diluted H_2_O_2_ was then directly added in BP/DMEM at a concentration of 1 mM for 1 minute. After 1 minute, the supernatant was collected and 15 mL of a quenching solution (10 mM sodium ascorbate – 5 mM Trolox – 10 mM sodium azide solution) was added to the dish to stop the biotinylation reaction.

Three washing steps were repeated in 20 mL quenching solution. Between each washing step, cells were centrifugated for 4 minutes. The quenching solution was then discarded. A centrifugation for 10 minutes was then performed. The quenching solution was aspirated and the dry pellet was lysed. RIPA lysis buffer supplemented with 1X protease inhibitor cocktail and with quencher at 1X was used. RIPA lysis buffer was made in a total volume of 50 mL, containing: 12.5 mL Tris Hcl (250 mM), 7.5 mL NaCl (750 mM), 1.25 mL SDS 20% (0.5% final), 12.5 mL NaDeoxy 10% (2.5% final),2.5 mL NP40 (5% final) and 13.75 mL H2O. For each condition, 500µL of lysis buffer was used. Samples were vortexed to facilitate lysis. Protein concentration was measured with Pierce 660nm protein assay kit (ThermoFicher scientific). 2 mg of proteins was used for enrichment on 100µL of streptavidin magnetic beads (ROCHE, ref 11641786001) at +4°C for 2 hours. The supernatant was then discarded and beads washed at 4°C for 5 minutes twice with RIPA buffer, once with KCl (1M), once with Na_2_CO_3_ (0.1M) and twice Urea (2M) in Tris HCl 10 mM at pH 8.0.

Proteins were then eluted for 10 minutes at 95°C in a protein loading buffer supplemented with biotin (2 mM) and DTT (20 mM). Samples were then quickly vortexed and cooled on ice for 5-10 minutes. Beads were then pelleted using a magnetic rack, frozen at -80°C and sent to the mass spectrometry (MS) platform EDyP in Grenoble on dry ice.

### Statistical analysis

If samples followed a Gaussian distribution, one-way ANOVA or two-way ANOVA were applied, either paired-ratio or unpaired depending on the experimental data. When samples did not pass the normality test, non-parametric test was applied (Mann-Whitney for unpaired samples and Wilcoxon signed-rank test for paired samples). * : p<0,05; ** : p<0,01; *** : p<0,001.

## ABBREVIATIONS

APEX: Ascorbate Peroxidase
ATG: Autophagy related genes
CNS: Central Nervous System
COP-II: Coat Protein Complex II
DKK1: Dickkopf-Wnt Signaling Pathway Inhibitor
DR: Dependence Receptor
ERGIC: ER-Golgi intermediate compartment
LRP5/6: LDL Receptor Related Protein 5
PLA: Proximity Ligation assay
STING: Stimulator Of Interferon Response CGAMP Interactor
TCGA: The Cancer Genome Atlas
WNT: Wingless-related integration site

## Notes

### Competing Interest Statement

The authors have declared no competing interest.

### Summary of Updates

The manuscript has been updated to include new experiments (primarily in the new Figure 4). The supplementary data have also been revised. The introduction has been simplified, and the discussion has been strengthened.

