## Supplemental data for "Kremen1 dependence receptor induces SEC24C and ATG9A-dependent autophagic cell death"

Supplementary Figure 1

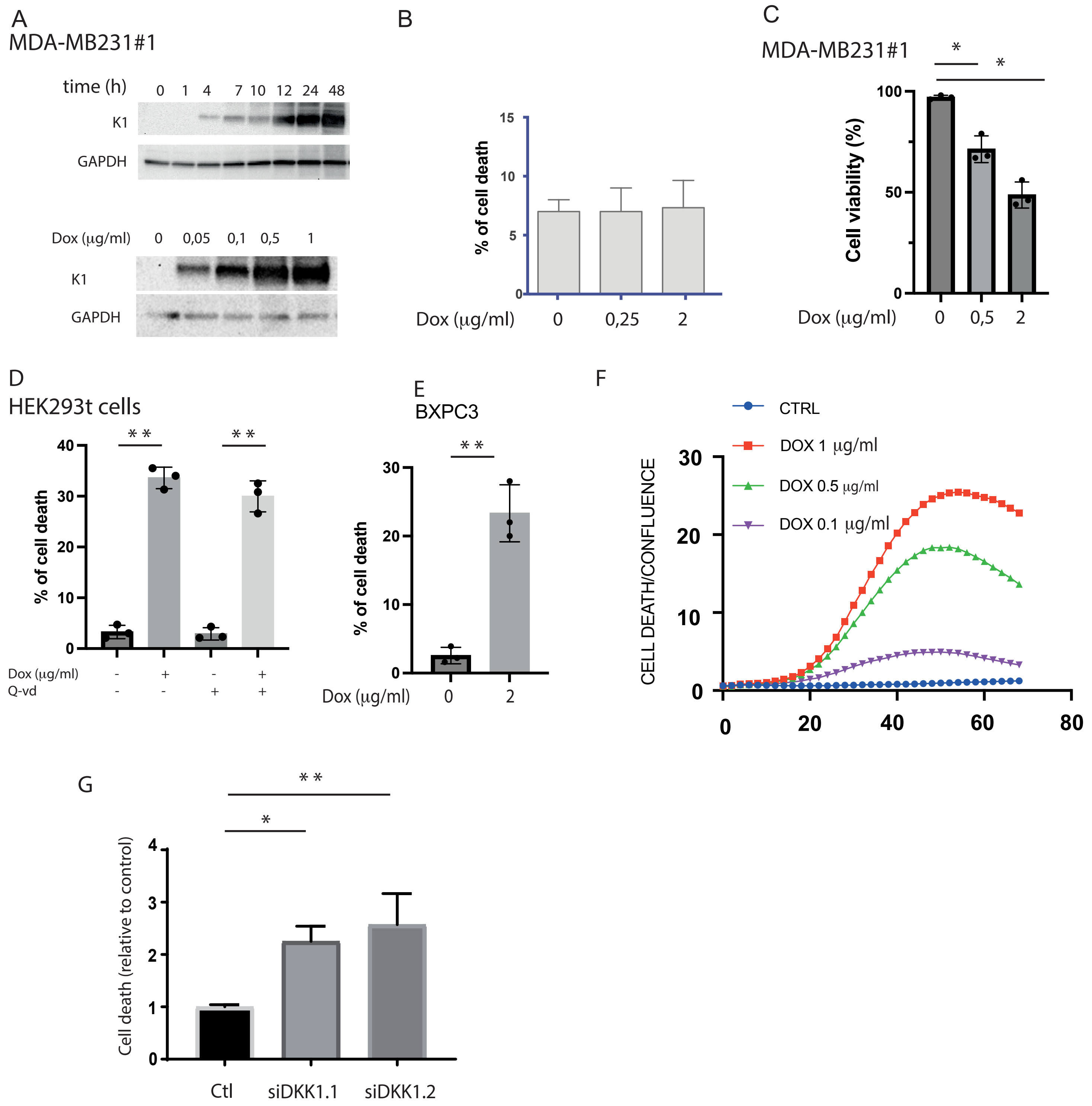


(A) Western blot analysis of Kremen1 expression in MDA-MB231K1 cells following doxycycline treatment after the indicated times and with the indicated concentration . (B), Cell death was assessed by trypan blue staining in cells expressing an empty vector following doxycycline treatment with the indicated concentrations. (C) Cell viability assessed by trypan blue staining in MDAMB231 cells expressing Kremen1 following doxycycline treatment with the indicated concentration at 48h. Paired-ratio Student's t-test was used with three independent experiments. (*: p-value<0.05 ; **: p-value<0.01). (D) Cell viability assessed by trypan blue staining in HEK293t cells expressing Kremen1 following doxycycline treatment with the indicated concentrations at 48h. Paired-ratio Student's t-test was used with three independent experiments. (*: p-value<0.05 ; **: p-value<0.01). (E) Cell viability assessed by trypan blue staining in BxPC3 cells expressing Kremen1 following doxycycline treatment with the indicated concentration at 48h. Paired-ratio Student's t-test was used with three independent experiments. (*: p-value<0.05 ; **: p-value<0.01). (F) Cell death was assessed in BxPC3 cells following induction of Kremen1 by doxycycline treatment with the indicated concentrations. Two way ANOVA wes used to test effect of Kremen1 induction in three independent experiments (**: p-value<0.01). (G) Cell death was assessed in MDAMB231 cells after 48h knocking down of DKK1 with two different siRNA. Paired-ratio Student's t-test was used with three independent experiments. (*: p-value<0.05 ; **: p-value<0.01).

Brahim Schott and Ghasemi supplementary figure 2

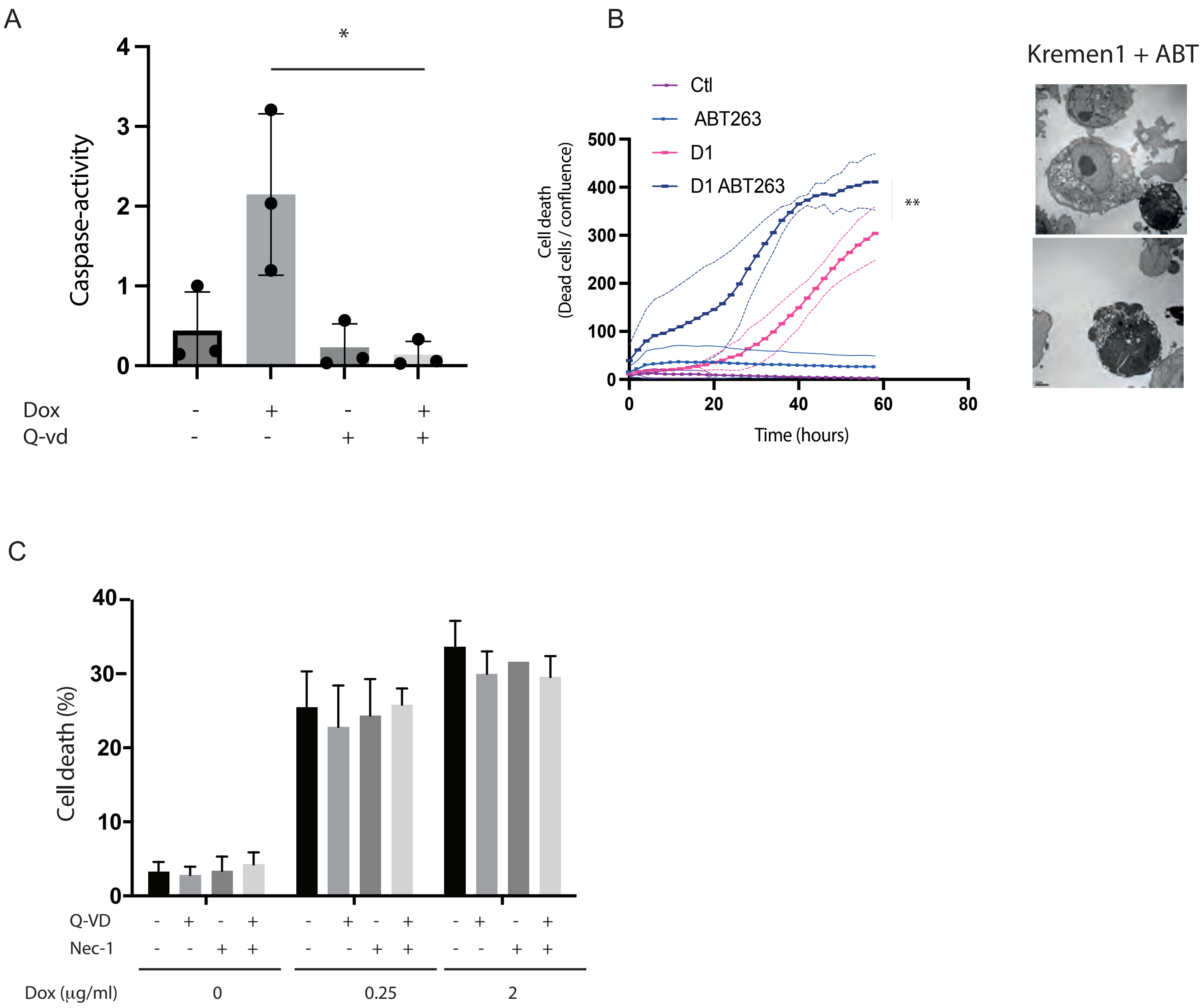


(A) Caspase-3 activity was measured in presence or not of Kremen1 expression (dox 1µg/ml) and q-VD-oph treatment (5µM). Paired ratio Student's t-test was used to assess the effect of q-VD in three independant experiments. (*; p-value < 0.05). (B) Incucyte monitoring of cell death of MDAMB 231 cells following doxycyclin treatment (D1; Dox 1µg/ml) in presence or not of ABT263 (1 µg/ml). Two way ANOVA was used to assess the effect of treatment on cell death (**: p-value < 0.01). (C) Cell death assessed by trypan blue staining in HEK293T cells treated with doxycycline at the indicated concentration in presence or not of Q-VD (10µM) or necrostatin-1 (10µM).

Phagophores Autophagosomes multivesicular bodies

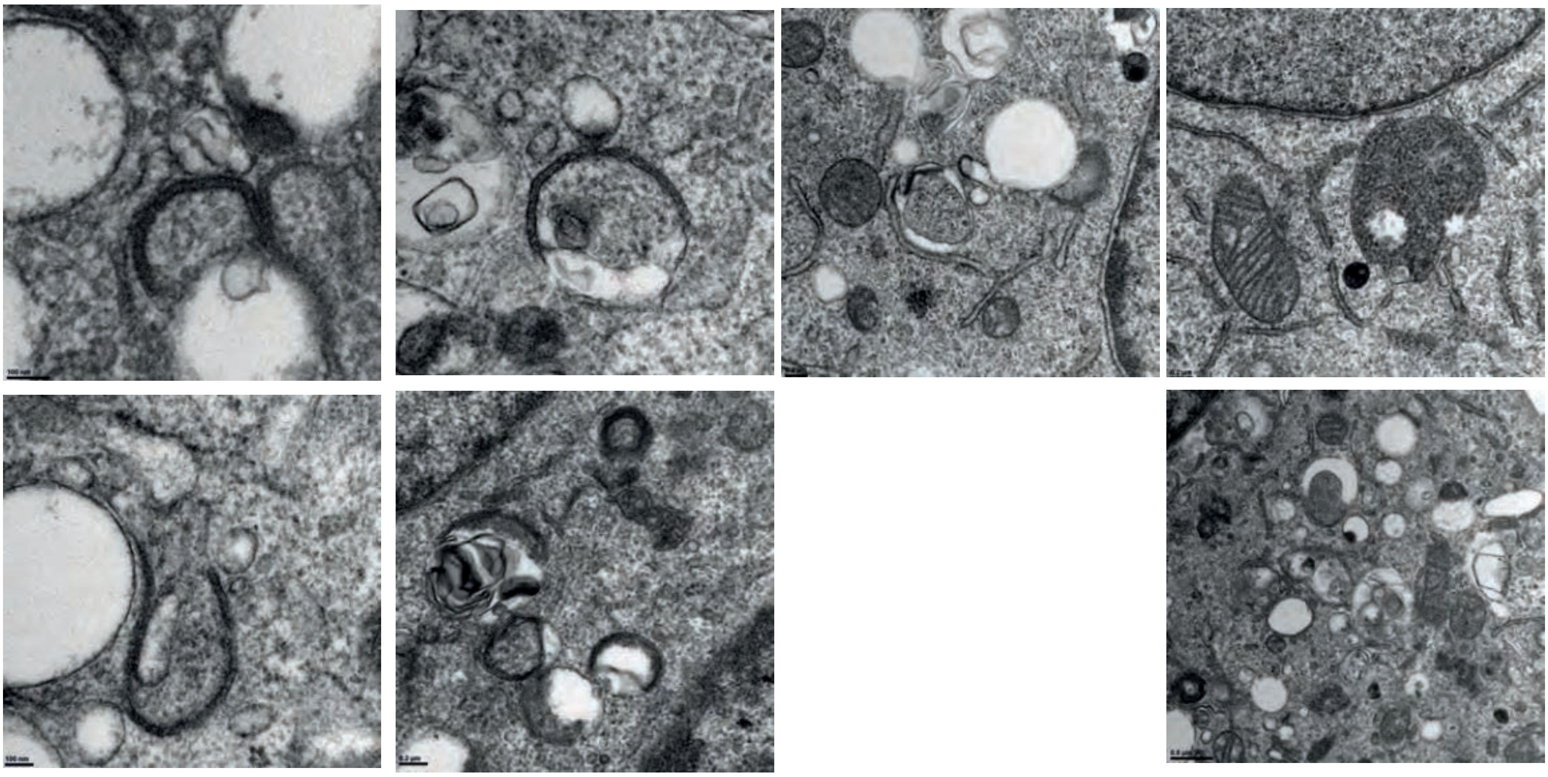

Supernatant

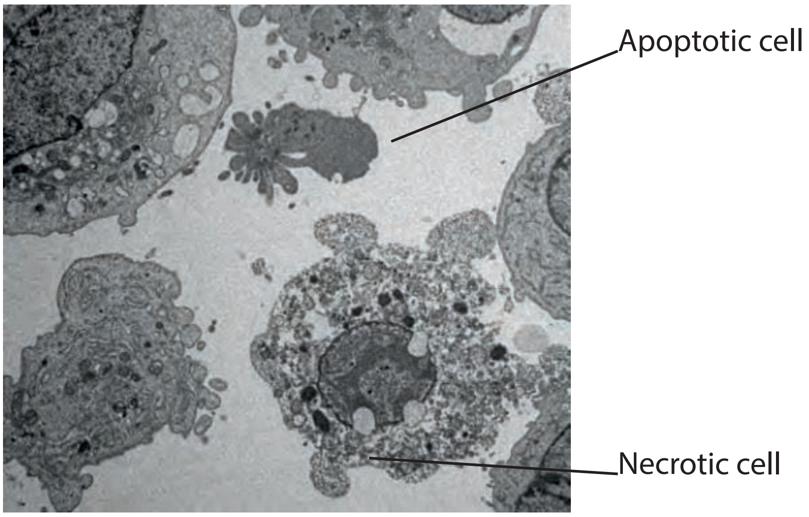


Supplementary figure 3. TEM observation of cells dying following Kremen1 induction. A Autophagic structures observed in adherent dying cells. B, Example of rarely observed apoptotic cells next to necrotic cells showing accumulation of vacuoles before permeabilization.

Supplementary Figure 4

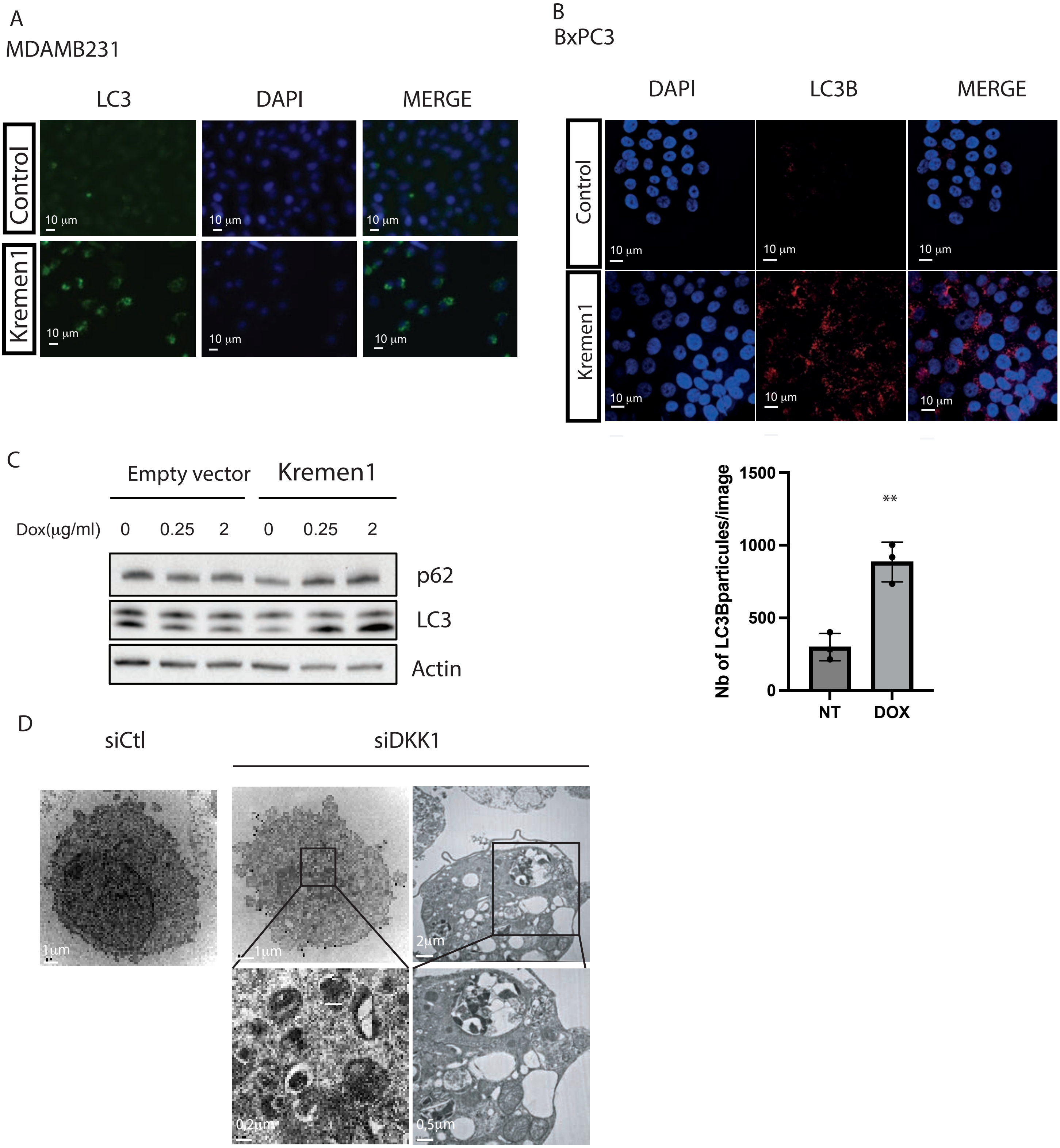


(A) Immunofluorence for LC3 in MDAMB231 cells treated (Kremen1) or not (Control) wiyth doxycycline (1µg/ml) for 24 hours. (B) Immunofluorence for LC3 in BxPC3 cells treated (Kremen1) or not (Control) wiyth doxycycline (1µg/ml) for 24 hours. (C) Western blot for p62 and LC3 in cell lysate of MDAMB231 treated with doxycycline to induce Kremen1. (D) TEM images of cell transfected with a control siRNA or a siRNA targeting DKK1.

Supplementary figure 5

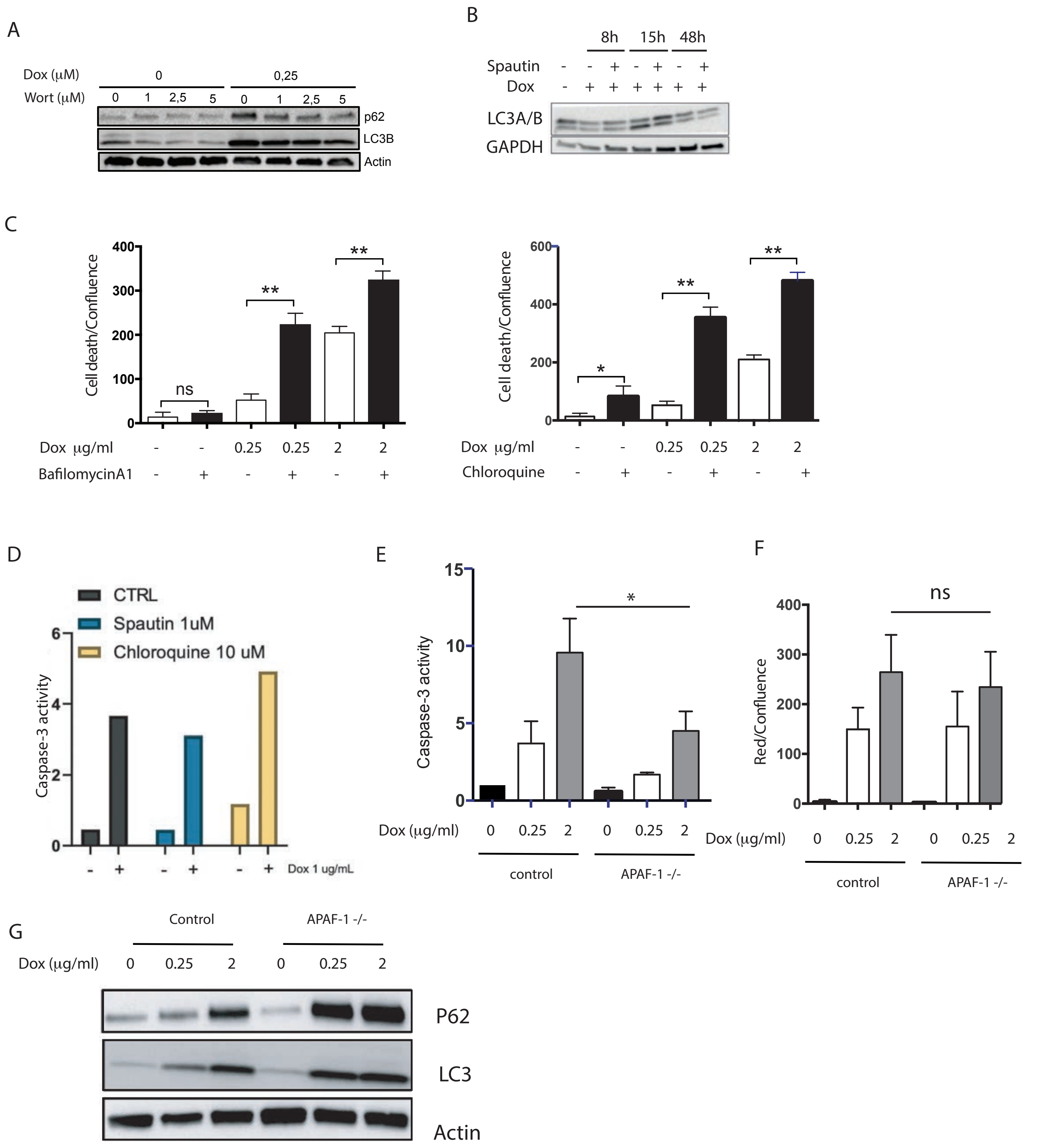


(A) Western blot analysis for LC3 and p62 following Kremen1 induction (dox, 0,25 μg/ml) for 24h, in presence of the specified concentration of wortmanin. (B) Western blot analysis for LC3 and p62 following Kremen1 induction (dox, 0,25 μg/ml) for 24h, in presence or not of Spautin1 (1μM). (C) Cell death was measured by incucyte in MDAMB231 following Kremen1 induction by doxycycline treatment with the indicated concentration in presence or not of bafilomycineA1 or Chloroquine. (D) Caspase 3 activity was measured following Kremen1 induction in presence or absence of Spautin and Choroquine. (E) Caspase-3 activity was measured in MDAMB231 or MDAMB231 cells knocked out for Apaf-1 (Apaf $1^{-/-}$) after treatment with doxycycline with the indicated concentration for 24 hours. (F) Cell death was measured in MDAMB231 or MDAMB231 cells knocked out for Apaf-1 (Apaf $1^{-/-}$) after treatment with doxycycline with the indicated concentration for 24 hours. Paired ratio student's t-test was performed on three independent experiment. *: p-value<0,05. (G) Western blot analysis of p62 and LC3 on cell lysates from WT MDAMB231 or MDAMB231 knocked-out for Apaf-1 after induction of Kremen1 with the indicated concentrations of doxycycline.

Supplementary Figure 6

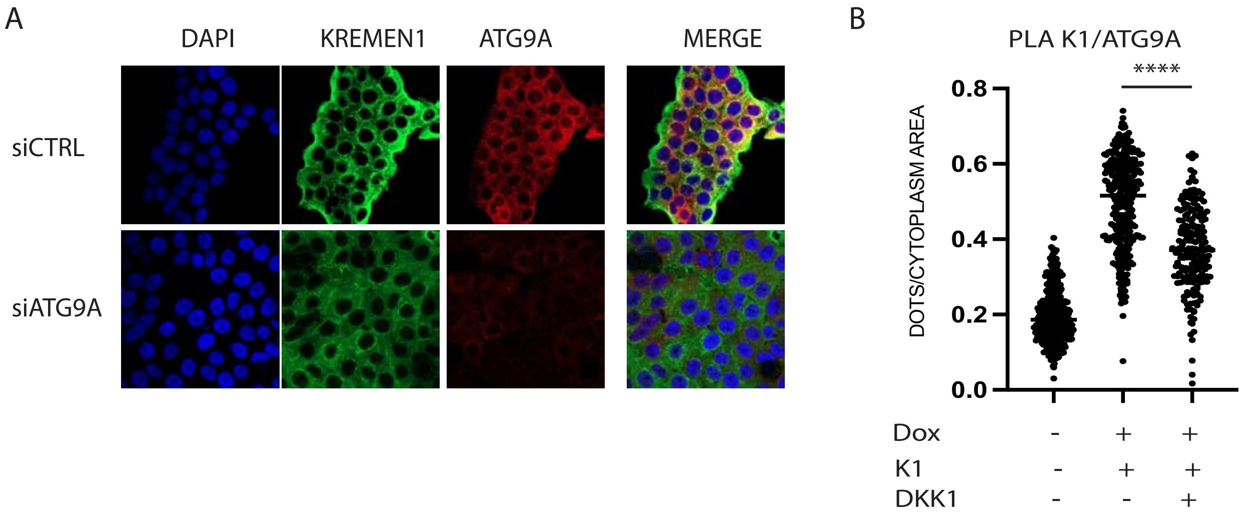


(A) Confocal analysis of cells stained with Kremen1, ATG9A in cells transfected with a siRNA control or a siRNA targeting ATG9A. (B) PLA analysis of the proximity of ATG9 and Kremen1 in presence or not of DKK1 expression. One way ANOVA followed by tukey's multiple comparison was performed. Results for comparison between DOX and DOX+DKK1 is specified. ****: p-value<0,0001.
